# The Mathematics Underlying Eeg Oscillations Propagation

**DOI:** 10.1101/2020.01.15.908178

**Authors:** Arturo Tozzi, Edward Bormashenko, Norbert Jausovec

## Abstract

Whenever one attempts to comb a hairy ball flat, there will always be at least one tuft of hair at one point on the ball. This seemingly worthless sentence is an informal description of the hairy ball theorem, an invaluable mathematical weapon that has been proven useful to describe a variety of physical/biological processes/phenomena in terms of topology, rather than classical cause/effect relationships. In this paper we will focus on the electrical brain field – electroencephalogram (EEG). As a starting point we consider the recently-raised observation that, when electromagnetic oscillations propagate with a spherical wave front, there must be at least one point where the electromagnetic field vanishes. We show how this description holds also for the electric waves produced by the brain and detectable by EEG. Once located these zero-points in EEG traces, we confirm that they are able to modify the electric wave fronts detectable in the brain. This sheds new light on the functional features of a nonlinear, metastable nervous system at the edge of chaos, based on the neuroscientific model of Operational Architectonics of brain-mind functioning. As an example of practical application of this theorem, we provide testable previsions, suggesting the proper location of transcranial magnetic stimulation’s coils to improve the clinical outcomes of drug-resistant epilepsy.

---

Spontaneous topological modifications in physical/biological systems may lead to novel functional features, indirectly dependent of exerted physical forces (Tozzi and Papo 2019). Here we consider one of the most intriguing theorems of algebraic topology, the “hairy ball” theorem (HBT, or Poincaré–Brouwer theorem), which states that there is no non-vanishing continuous tangent vector field on even-dimensional *n*-spheres (Milnor 1978; Eisenberg and Guy, 1979). A naïve description asserts that “given at least some wind on Earth, there must at all times be a cyclone or anticyclone somewhere”. HBT ensures the presence of at least one point on the sphere where the tangential components of vectors (and/or tensors) disappear. HBT relates several physical phenomena to non-local topological effects, rather than to local physics (Tozzi and Papo, 2019; Bormashenko 2016). Indeed, HBT has been used to analyze nematic solid shells deformations (Modes and Warner, 2012), nanoparticle chains growth (DeVries et al, 2007), flux line patterns occurring when a magnetic field is applied to a type-II superconducting crystal (Layer and Forgan, 2010), patterns arising from dynamic surface - Marangoni-like- instabilities (Bormashenko 2015) and the spin-base invariant formalism of Dirac fermions (Giesand Lippoldt, 2015). Novel feasible applications of HBT have been recently suggested (Tozzi and Papo 2019) for the assessment of cell membrane surface tension, Uhlenbeck’s singularity removal theorem (Uhlenbeck, 1982; Tao and Tian 2004), dewetting transitions (Sharma and Reiter, 1996; Li et al., 2019), unconventional superconductors (Laver and Forgan, 2010), surface instabilities inherent for liquid/vapor interfaces (Bormashenko 2015), non-Hermitian degeneracies (Miri and Alù, 2019) which naturally arise in sparse neural networks (Amir et al., 2016). HBT describes the collective dynamics of particles in spherical crystals (Yao, 2019): in particular, it predicts defect-driven synchronized breathing modes, emerging around disclinations related to disruption of crystalline order (Yao, 2019). Furthermore, HBT zero-velocity points appear in numerous physical problems, including rolling of solid bodies (Bormashenko and Kazachkov, 2017) and Leidenfrost droplets (Bormashenko 2019).

Here we start from the recent observation that HBT is also able to describe electromagnetic waves propagation (Bormashenko 2016). When oscillations in a given medium display wave fronts topologically equivalent to a sphere, there must be at least one point of the wave front where electromagnetic fields’ vectors, speed and Poynting vector equal zero. There exist certain so-called “dark directions”, well-known to engineers designing antennas (Liang et al. 2013), devoid of energy transport (Chen et al, 2019) and unavoidable for electromagnetic fields generated by multipoles characterized by vector spherical harmonics (Chen et al., 2019). Here we make an effort to describe the restrictions imposed by HBT on the spatial distribution of vectors 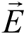 in the propagating electric waves extracted from electroencephalogram (EEG) traces. Provided that the electric waves propagate spherically on the cerebral cortex, HBT dictates the occurrence of at least one point on the cortical surface at which vectors of the electric fields equal zero. In other words, when a continuous tangential velocity field exists on a surface homeomorphic to a ball, zero velocity points will be necessarily present at the surface. We show how this observation has major implications for the assessment of brain metastability, as well as for clinical applications: for example, the search for target zones for the use of transcranial magnetic stimulation (TMS).

## MATERIALS AND METHODS

### Hairy ball theorem and wave fronts propagation in physical systems

Consider propagation of electromagnetic fields, when the wave front (i.e. the surface of the constant phase) displays a shape that is topologically equivalent to a sphere (i.e., the surface is characterized by the Euler number *χ* = 2). In this case, the vectors of the electric 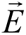 and magnetic 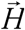 fields form continuous tangential fields (**Figure 1A)**. HBT demands that the tangential component of the continuous vector field, defined on the surface topologically equivalent to a sphere, will be zero in at least one point located on the surface. For further details, see Liang et al. 2013, Bormashenko, 2016, Chen et al, 2019. Therefore, according to HBT, both 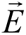 and 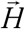 have at least one point where fields are zero. When both the fields simultaneously equal zero in the same point, we achieve 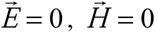, due to the equivalence 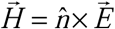, where 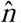 is the unit vector normal to the wave front. In the same point, the Poynting vector 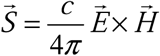 equals zero too. Because the group velocity of electromagnetic wave in this direction is zero, no transport of energy in the direction normal to the wave front in this point is possible and the “dark direction” is formed, whatever the physical source field could be (Liang et al. 2013, Chen et al, 2019). To provide an example, consider the hypothetical field (either electromagnetic or hydrodynamic, and so on) in **Figure 1B**, where the wave progression of the vectors displays a positive-curvature front. According to the HBT dictates, a point with zero value must exist on the wave front (**Figure 1C**), which causes a local perturbation of the field and leads to wave front deformation (**Figure 1D**).

**Figure 1.**
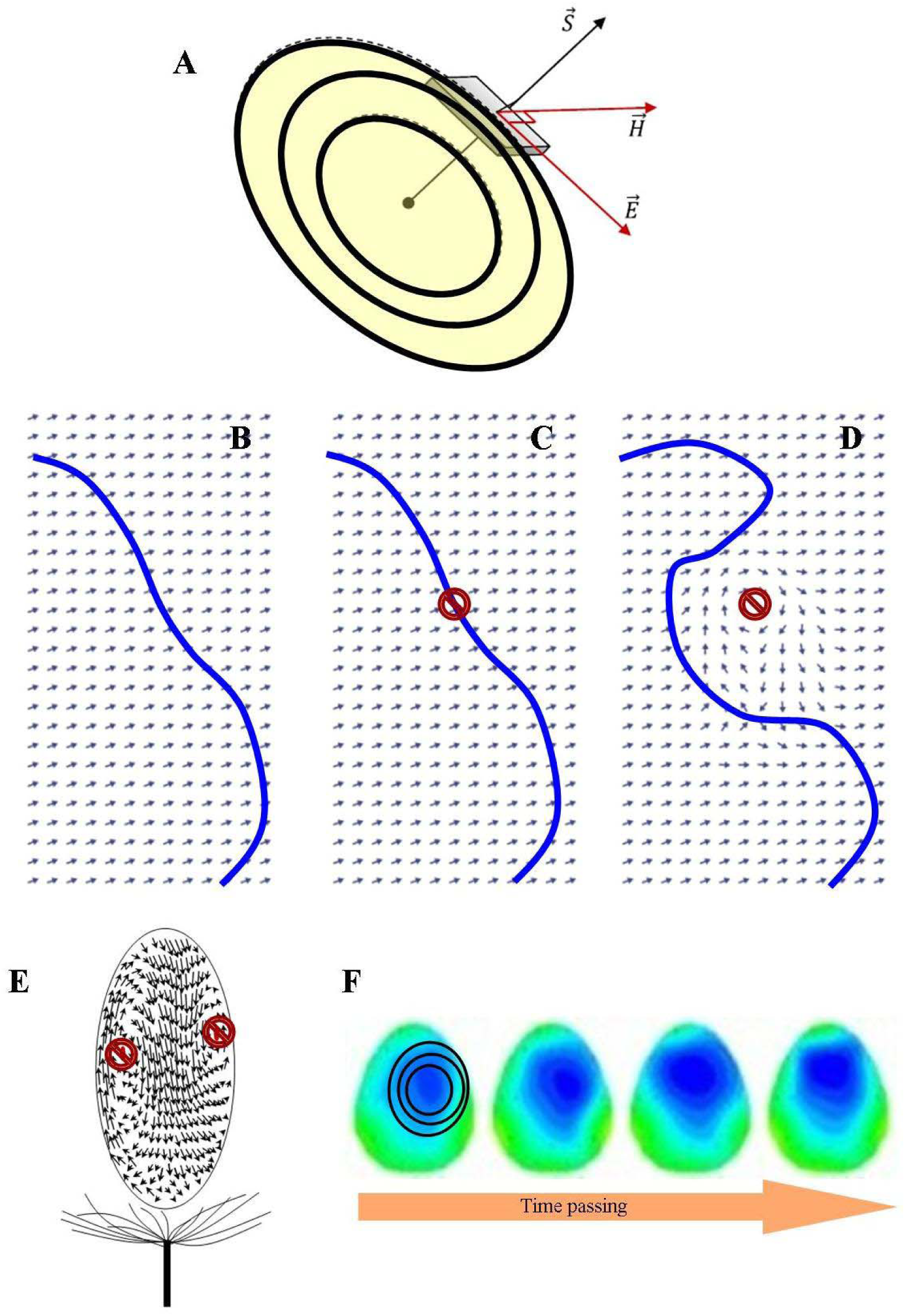
Schematic illustration of pattern propagation of oscillations in physical systems, including the brain. **Figure 1A**. Propagating electromagnetic wave. 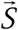 stands for the Poynting vector. Note that the concentric patterns of electric propagation give rise to a wave front with a surface topologically equivalent to a sphere. Modified from Bormashenko (2016). **Figures 1B-D**. Simulation of a hypothetical world sheet in spacetime depicting a generic flow equipped with a positive-curvature wave front (modified from Beekman, 2011). When the wave front proceeds (**1B**), the occurrence of the tangential components of vectors (**1C**) leads to modifications in vectors trajectories and, consequently, to distortions in the wave front (**1D**). In this simulation, the vanishing point of the wave front predicted by HBT is illustrated by a prohibition sign. **Figure 1E**. Sketch of dandelion and the vortexes it generates. The arrows illustrate the field of velocities at the vortex surface. The two prohibition signs depict the stagnation points dictated by HBT. Modified from Cummins et al. (2018). **Figure 1F**. The concentric, positive-curvature patterns typical of electromagnetic fields can be also found when examining the electric waves’ temporal propagation in two-dimensional EEG traces.

The vector field in principle may be normal to the surface in this point. Apparently it is impossible for the vectors of the electric and magnetic field to be normal to the surface of the constant phase (in other words parallel to the wave vector), due to the transverse nature of the electromagnetic field. It is noteworthy that the vectors of electric and magnetic fields will possess a zero point placed on the wave front at any given time, if the surface of the constant phase keeps the shape topologically equivalent to a sphere. It is also worth to be underlined that HBT holds not just for spherical manifolds, but also for every manifold with positive-curvature, provided its genus is zero. For example, Cummins et al. (2018) analyzed the motion of wind dispersed plants, reporting a vortex of recirculating air, which is detaching owing the flow passing through the pappus (**Figure 1E**). The occurrence of stagnation points (in this case two instead of one) can be noticed where the air velocity at the vortex surface is zero. The dandelion example suggests that the manifold does not need necessarily to be a sphere: HBT just requires that such manifold should be topologically equivalent to a sphere. This observation will have important implications in the following.

### Looking for HBT in nervous electric waves

In the sequel, we provide an effort to use HBT to analyze the EEG electric activity of the brain. We choose to focus on the ELECTRIC 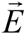 field of the brain, rather than the MAGNETIC field 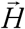, because the former is more powerful and easier to detect through simple EEG techniques. Indeed, the circular patterns and wave fronts of electric waves propagation can be easily identified in real EEG traces. EEG traces do encompass electric waves with sphere-like wave fronts (**Figure 1F**). The requirement of spherical currents is satisfied when we analyze brain electric activity: indeed, the shape of flux surfaces that contains current lines is closed and orientable in the brain, so that it is homeomorphic to a sphere *S*^2^. Since the electric field in the brain is nowhere vanishing and tangent to the surface, HBT rules out the possibility to describe nervous oscillations as taking place on a sphere. Also, the “dark directions” can be predicted for the electric field generated by brain, when the field appears on the cortical surface topologically equivalent to a sphere.

According to HBT, the circular propagation of concentric electric waves must display zero-points on their wave fronts. Further, when electric waves propagate from a given brain area, the wave front must necessarily be locally perturbed by these zero-points. For the purpose of this study to assess whether and how HBT permits a novel description of electric fields’ propagation in the brain, we analyzed EEG traces of three subjects that were presented emotional visual stimuli, according to the procedure described by Jaušovec and Jaušovec (2010). The emotional stimuli were color slides selected from the International Affective Pictures System (Lang et al., 2005) according to the valence dimension: emotionally positive (valence ratings from 7.2 to 8.2), neutral (valence ratings from 4.4 to 6.2), and negative (valence ratings from 1.3 to 2.0). The discrimination of picture categories occurred during passive viewing and was internally driven – the respondents were not asked to make any judgments or motor responses. EEG was recorded using a Quick-Cap with sintered (Silver/Silver Chloride; 8mm diameter) electrodes. Using the Ten-twenty Electrode Placement System of the International Federation, the EEG activity was monitored over nineteen scalp locations (Fp1, Fp2, F3, F4, F7, F8, T3,T4, T5, T6, C3, C4, P3, P4, O1, O2, Fz, Cz and Pz) during the visual task. ll leads were referenced to linked mastoids (A1 and A2), and a ground electrode was applied to the forehead. Additionally, vertical eye movements were recorded with electrodes placed above and below the left eye. The digital EEG data acquisition and analysis system (SynAmps) had a bandpass of 0.15-100.0 Hz. At cutoff frequencies, the voltage gain was approximately –6dB. The 19 EEG traces of the three subjects were digitized online at 1000 Hz with a gain of 1000 (resolution of 084 μV/bit in a 16 bit A to D conversion), and stored on a hard disk. Epochs were automatically screened for artifacts. All epochs showing amplitudes above +/-50 microV (less than 3%) were excluded, to avoid that the traces could be artifacts of the visualizing algorithm while plotting the EEG power maps. The EEG study was done according with Declaration of Helsinki and was approved by the Ethics Committee of the University of Maribor, Slovenia.

To practically identify the occurrence of zero-points of electric activity in the real EEG traces, we evaluated two different plots: a) the raw data, i.e., a table displaying the electric values in μV in every cortical area for every time in ms; 2) a 2-D temporal reconstruction of EEG waves. We evaluated the changes in electric activity occurring in the first 500 ms after the presentation of emotional stimuli.

## RESULTS

According to our topological approach, the concentric wave fronts of EEG electric activity can be assessed in terms of HBT. When EEG traces display zero values in some areas at a given time, predictable changes in electric wave fronts must occur elsewhere. Looking at the table of EEGs raw data from the three subjects presented with visual stimuli, we detected the occurrence of about 15-20 zero-points/500 ms, with different electrode scalp locations and different timing (**Figure 2**). We did not find a predictable path in both timing and location of the newly-formed zero-points.

**Figure 2.**
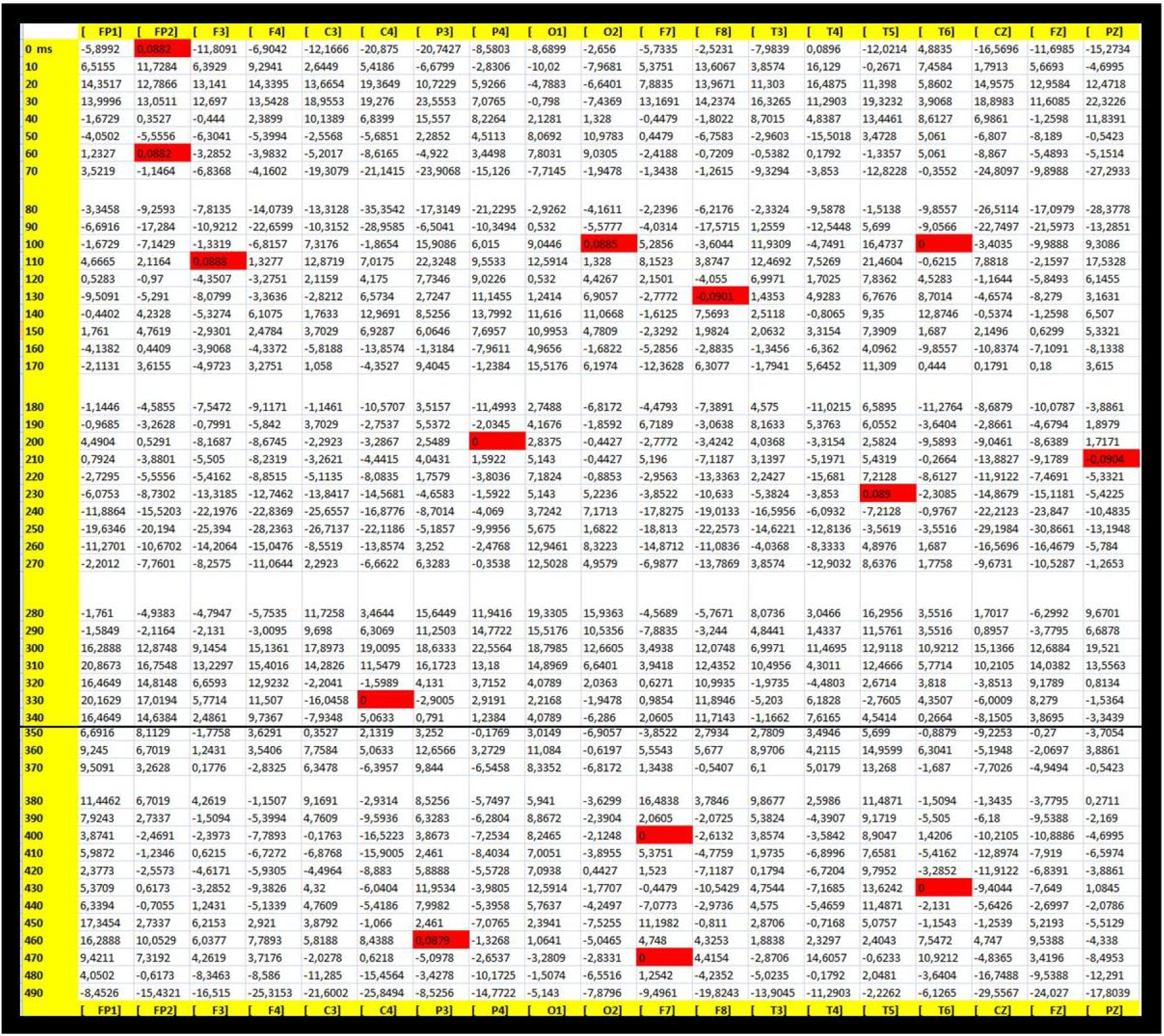
Raw data from EEG activity in a subject who underwent emotional stimuli. The results are plotted as electric activity (in μV) across different brain areas vs time locations (in ms). The red squares illustrate the electrode location and the timing of the electric activity almost equaling zero μV.

To compare the occurrence of zero-points in raw data with the modifications of electric wave fronts, we examined the corresponding EEG 2D temporal plots from the three subjects (**Figure 3A**). A spatio-temporal correlation can be found between the sudden occurrence of zero-points and the subsequent onset of warped wave fronts. Indeed, almost ubiquitously, the occurrence of a zero-point is followed with a delay of 20-30 ms by the formation of a wave front in areas far apart the zero-points. See **Figure 3B** for further details.

**Figure 3.**
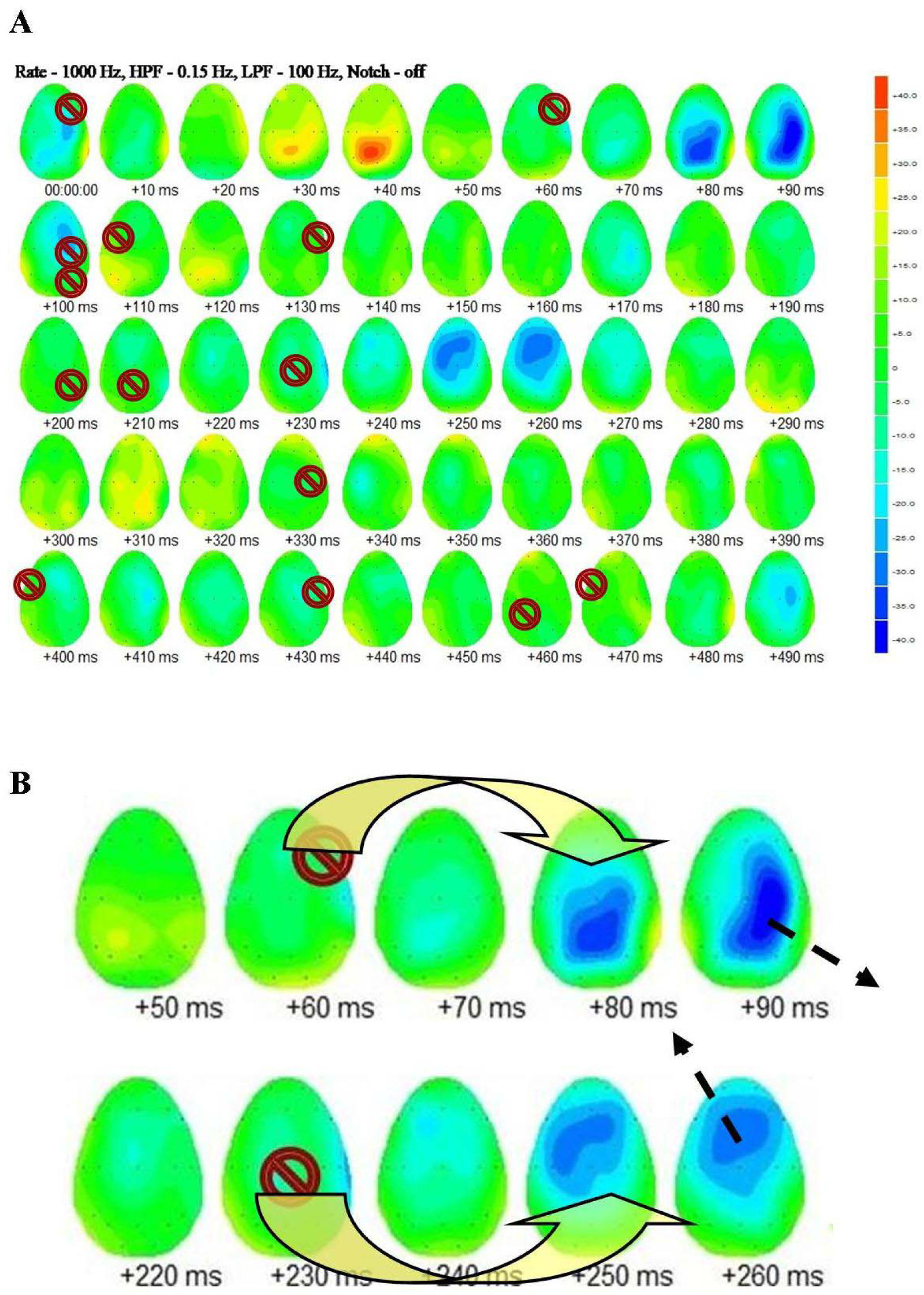
The role of HBT in modifying the electric activity of the brain in a subject presented with visual stimuli. The prohibition signs illustrate the points where electric zero-values occur in the first 500 ms after the stimuli. Note the occurrence of wave fronts, depicted as concentric increases or decreases of cool and warm colors. **Figure 3B**. Magnification of two short frames from the Figure 3A. When the vanishing zero-points (prohibition signs) appear in a given area, concentric wave fronts (blue concentric areas) arise in different areas after about 20-30 ms (curved arrows) and tend to propagate in a direction opposite to the zero-points (straight dotted arrows). Therefore, the occurrence of vanishing zero-points allows to predict the rise and the progression of the ensuing electric wave front.

In particular, in **Figure 3A**, the 14 detected zero-points were followed by the above-described, predictable electric wave front pattern in 13/14 (92.85%) cases.

Therefore, the occurrence of zero-points is able to modify the circular propagation of electric waves and to give rise to deformations of the wave fronts. In the case of the electric activity of the brain during visual stimulation, the occurrence of wave front modifications can be visually predicted when examining the location and the timing of the zero-points in the EEG two-dimensional plots.

## DISCUSSION

Starting from the observation that HBT constraints the vectors’ spatial distribution of electric and magnetic fields in the far Fraunhofer region (Bormashenko 2016), we used this simple theorem from algebraic topology to describe the electric activity (EEG) of the brain. Indeed, the vectors of electric fields are equipped at least with one zero point placed on the wave front at any given time, in case of the wave front’s shape is topologically equivalent to a sphere. In this study, we showed that zero-points can be reliably identified in EEG signals and that they give rise to predictable wave fronts of electric diffusion. In what follows we will place our findings in the context of brain-mind research, as well as provide an illustrative example for a clinical application.

One of the most promising theoretical neural frameworks is the general theory of brain-mind operational architectonics (OA) (Fingelkurts and Fingelkurts, 2004). According to OA, the simplest mental/cognitive operations are correlated with local 3D-fields produced by transient functional neuronal assemblies, while complex mental/cognitive operations are produced by joining a number of simple operations (temporal coupling of local 3D-fields by means of *operational synchrony*, OS) in form of metastable *operational modules* (OM) of varied life-span. Nested OA displays a peculiar brain *operational space–time* (OST) (**Figure 4B**) that is best captured by EEG measurement (Fingelkurts and Fingelkurts, 2017). At the EEG level, simple mental operations (phenomenal qualities, emotions, and so on) are equivalent to the EEG quasi-stationary segments, within which the local fields generated by transient functional neuronal assemblies are expressed. The quasi-stationary EEG segments within each local EEG signal are marked by boundaries in the form of *rapid transitional processes/periods* (RTPs) (**Figure 4A**), i.e., abrupt EEG amplitude changes observed within a short-time window. We suggest that RTPs could be correlated with the occurrence of HBT zero-points. Indeed, in comparison to the length of quasi-stationary segments, each RTP has a very short duration and can therefore be treated as a point, i.e., the HBT zero-point of our topological approach. Furthermore, the number of RTPs fits well with the number of HBT zero points: the number of RPTs per each minute local-EEG (at least during restful wakefulness with closed eyes) is about 61-351, depending on the frequency range, with a progressive increase from delta to gamma frequency range (Fingelkurts and Fingelkurts, 2015). In turn, the RTP range of temporal durations is produced on timescales corresponding to 100-900 ms range, while the occurrence of zero-points is followed by changes in electric wave fronts after 20-30 ms.

**Figure 4.**
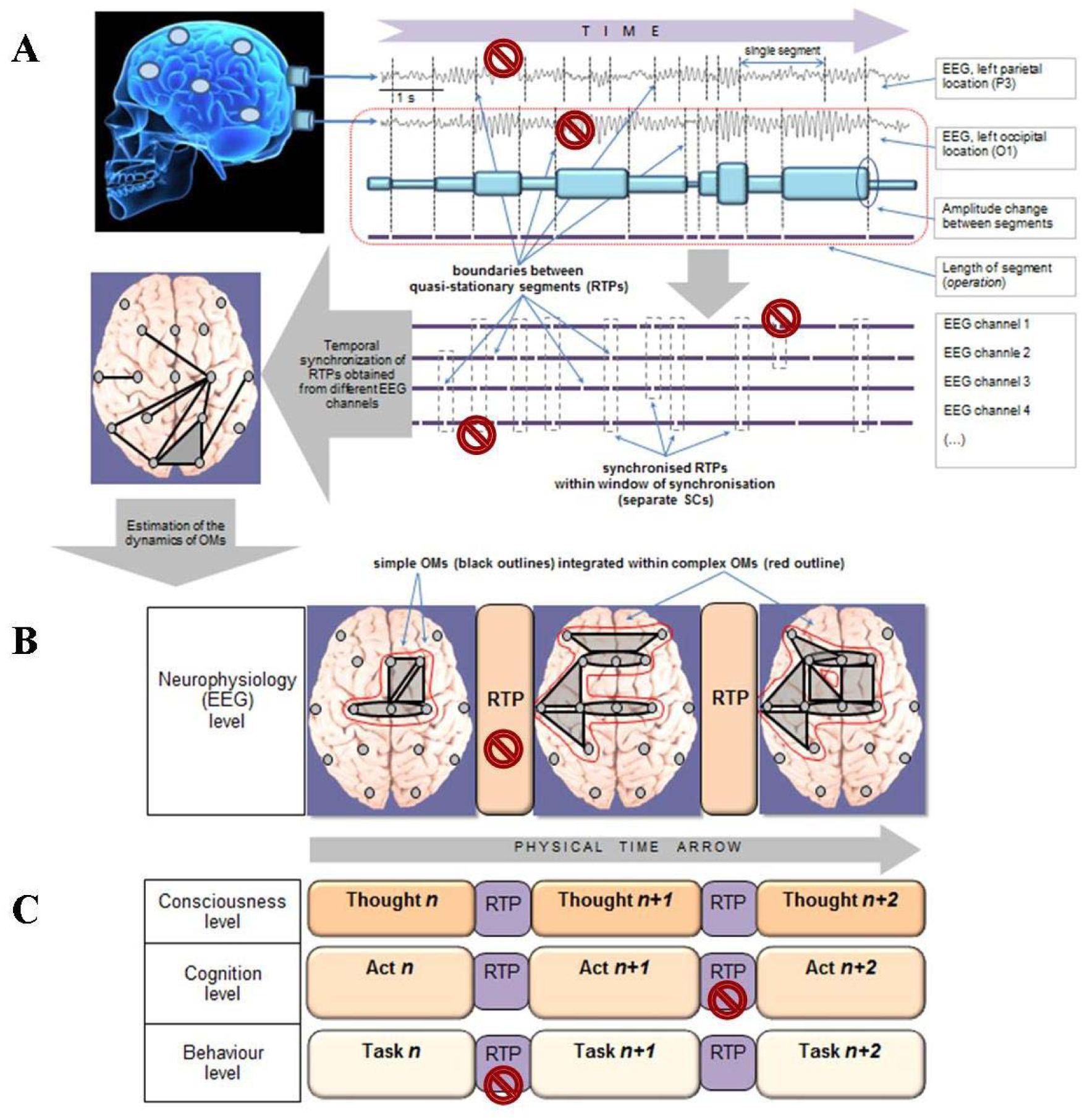
Schematic illustration of the Operational Architectonics methodology and possible relation to HBT. All RTPs should be HBT zero-points: here, for sake of simplicity, just a few HBT zero-points are illustrated (prohibition signs). It must be reminded, however, that all RTPs should be HBT zero-points. **Figures 4A-C**. Schematic illustration of EEG assessment of (**A**) neuronal assembly’s dynamics and relation of this dynamics to simple operations and (**B**) nested large-scale conglomerates of synchronized neuronal assemblies in the form of nested operational modules (OMs) and (**C**) their relation to a stream of complex operations or cognitive/conscious acts. RTP – rapid transitional period (boundary between quasi-stationary EEG segments within the same local EEG signal); SC – synchrocomplex – momentary synchronization of RTPs among several local EEG signals within short temporal window of synchronization; Gray shapes illustrate individual (simple)OMs; Red line illustrates complex OMs. **Figure 4C** schematically depicts the functional structures of phenomenological consciousness, cognition and behavior functionally isomorphic with the structure of electrophysiological level. Cognitive, phenomenological, and behavioral levels illustrate the ever-changing stream of cognitive/phenomenal/behavioral acts, where each momentarily stable pattern is a particular cognitive/phenomenal/behavioral macro-operation (thought/image/act) separated by transitive fringes (RTPs). This figure is modified from Tozzi et al. (2017).

The transition from one segment to another reflects the moment of abrupt switching from one neuronal assembly’s operation to another (**Figure 2A**). The events similar to RTPs are referred to as renewal (or critical) events: namely, the crucial events that reset the memory of the system. This property is correlated with the physical phenomenon of “intermittency” and is compatible with self-organized criticality found both in physical systems and in the brain. Since the beginning and the end of discrete operations performed by local neuronal assemblies are marked by sharp changes (RTPs) in the amplitude of local EEG signals, the simultaneous occurrence of such RTPs from different local EEG signals within the multichannel EEG recording could provide evidence of synchronization of simple operations performed by neuronal assemblies (located in different brain areas) that participate in the same functional act as a group (**Figure 4B**), *e.g*., executing a particular complex operation responsible for a subjective presentation of complex objects, scenes or thoughts. HBT can be correlated with the general theory of brain-mind OA, that considers the brain as a metastable system (Fingelkurts and Fingelkurts, 2004, 2017): in HBT terms, the complex mental operations (reflected in the transitory spatiotemporal EEG patterns formed by synchronized quasi-stationary segments separated by the RTPs) could be correlated with the HBT-related electric zero-points that give rise to RTPs (**Figure 3C**).

One established the occurrence of events dictated by HBT, the next question is: has this observation testable applications? Below we illustrate one of the potential clinical applications of HBT. Let us consider the case of epilepsy. The occurrence of points where brain oscillations vanish provides us with a novel way to use (TMS). TMS is a focal electrical brain stimulation induced by powerful magnetic fields (Chung et al., 2016; Kohli and Casson, 2019). Low frequency repetitive rTMS (0.3-1 Hz) decreases cortical excitability, suggesting a potential therapeutic advantage for patients with drug-resistant epilepsy (Jan et al., 2017; Gersner et al., 2016). Chen et al. (2016) retrospectively analyzed the evidence for the efficacy of TMS in drug-resistant epilepsy, evaluating studies that used rTMS of any frequency, duration, intensity and setup (focal or vertex treatment). Excruciatingly, the evidence for efficacy of rTMS for reduction in seizure rate/frequency is still lacking, due to the extreme variability in outcome reporting (Jan et al., 2017). There is still no agreement on optimized stimulation parameters and patterns of rTMS for epilepsy, because TMS effects vary across individuals and depend on a large number of factors: frequency, number of stimuli within a train, stimulation intensity, type of coil, coil position, duration of stimulation, and inter-train interval.

In this uncertain context, an HBT approach to epileptic seizures suggests that the area of onset of pathological spikes might be characterized by an impaired amount of topologically vanishing points: in particular, it might be hypothesized an increase in number of zero points (**Figure 5A**). The occurrence of vanishing points gives us the possibility to draw a testable prevision: to achieve the best therapeutic effect in epilepsy, rTMS treatments must use not solid, coherent beams, rather hollow magnetic beams, i.e., cylindrical beams with internal cavities devoid of magnetic stream. Indeed, the use of cylindrical hollow beams is suggested by HBT-related topology: when a sphere with genus zero becomes a torus (i.e., a manifold with genus equal or higher than one), HBT does not hold anymore. To provide an example, the **Figure 5B, left side**, illustrates the rotation of a rigid ball around a fixed axis OO_1_: it is easy to see that O and O_1_ are zero velocity points, in touch with HBT’s dictates. In turn, the zero velocity points disappear when the ball is drilled completely through, because HBT does not hold anymore in case of a manifold being a torus with genus one (**Figure 5C, left side**). The same holds for the circular wave fronts of the brain electric activity (**Figures 5B-C, right side**): in rTMS, terms, the hollow beam corresponds to the production of a torus inside the circular wave front (**Figure 5C, right side**). In case of a manifold with genus one artificially provoked by rTMS’s external waves, HBT does not hold anymore: this means that the normal electric activity of the brain with impaired zero-points can be restored. In a HBT account, epilepsy can be compared to Marangoni-like instabilities: the pathological zero-points are modified by rTMS-mediated forces, reversing the unwanted seizure oscillations and restoring the normal cortical function.

**Figure 5A.**
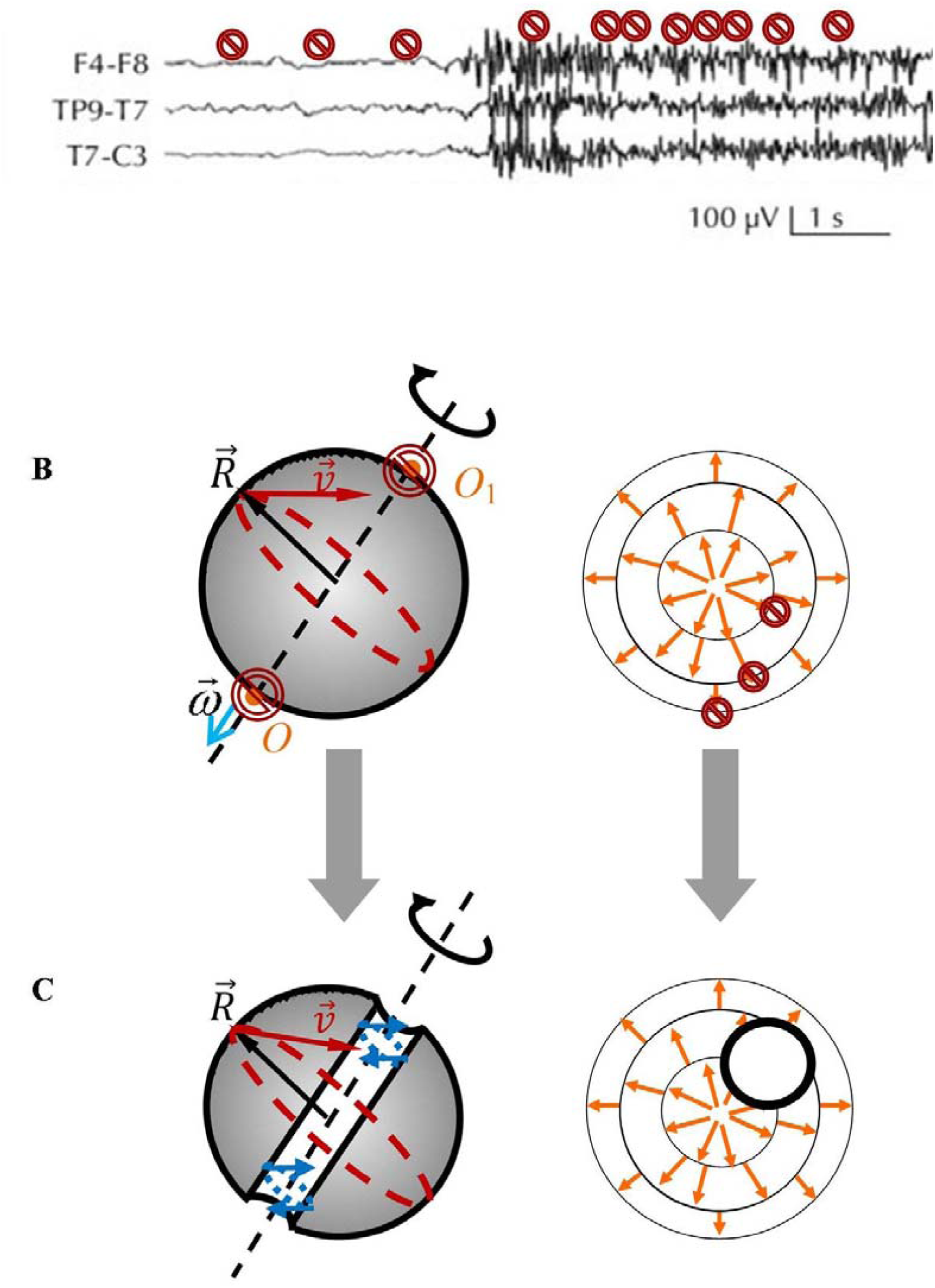
A Recorded surface EEG (displayed in a bipolar transverse montage) showing a left parietal EEG seizure onset. Modified from: Beleza et al. (2010). The number of vanishing points (prohibition signs) is depicted as increasing after the seizure onset. **Figure 4B**. Rotation of a rigid ball (left) compared with an electric concentric wave front (right). In both cases, the occurrence of zero-points (prohibition signs) is required by HBT. Modified from: Bormashenko and Kazachkov (2017). **Figure 4C**. The occurrence of a torus removes the zero-points in both the rigid ball and the electric wave front. See text for further details.

To sum up, clear-cut topological theorems, such as, e.g., the Borsuk-Ulam theorem described by Tozzi et al. (2017), can be used to make sense of experimentally observed neuroscientific phenomena. In particular, here we discussed the implications of HBT-correlated zero-valued points in nervous dynamical systems, describing how mathematical-related events, such as vector modifications, generate changes in brain features that comply with strict topological requirements.

## REFERENCES

1) Amir A, Hatano N, Nelson DR. 2016. Non-Hermitian localization in biological networks. Phys. Rev. E 93, 042310. doi: https://doi.org/10.1103/PhysRevE.93.042310.

2) Beekman A.J. 2011. Vortex Duality in Higher Dimensions. Casimir PhD Series. urn:isbn:9789085931133.

3) Beleza P, Rémi J, Feddersen B, Peraud A, Noachtar S. 2010. Epidural and foramen-ovale electrodes in the diagnostic evaluation of patients considered for epilepsy surgery. Epileptic Disord. 2010 Mar;12(1):48–53. doi: 10.1684/epd.2010.0297.

4) Bormashenko E. 2016. Obstructions Imposed by the Poincaré–Brouwer (“Hairy Ball”) Theorem on the Propagation of Electromagnetic waves. Journal of Electromagnetic Waves and Applications, 30(8). doi: /10.1080/09205071.2016.1169226.

5) Bormashenko E. 2015. Surface instabilities and patterning at liquid/vapor interfaces: Exemplifications of the “hairy ball theorem”. Colloids and Interface Science Communications. Volume 5, March 2015, Pages 5–7. https://doi.org/10.1016/j.colcom.2015.04.003.

6) Bormashenko E, Kazachkov A. 2017. Rotating and rolling rigid bodies and the “hairy ball” theorem, Am. J. Physics 85, 447.

7) Bormashenko E. 2019. Motion of the liquid on the surface of Leidenfrost droplets and the hairy ball theorem, Surf. Innovations 7 (2), 101–103.

8) Chen R, Spencer DC, Weston J, Nolan SJ. 2016. Transcranial magnetic stimulation for the treatment of epilepsy. Cochrane Database Syst Rev, (8):CD011025. doi: 10.1002/14651858.CD011025.pub2.

9) Chen W, Chen Y, Liu W. 2019. Singularities and Poincaré Indices of Electromagnetic Multipoles, Phys. Rev. Lett. 122, 153907.

10) Chung SW, Sullivan C, Rogasch NC, Hoy KE, Bailey NW, et al. 2019. The effects of individualised intermittent theta burst stimulation in the prefrontal cortex: A TMS-EEG study. Hum Brain Mapp.;40(2):608–627. doi: 10.1002/hbm.24398.

11) Court JH., Raven J. 1995. Manual for Raven’s progressive matrices and vocabulary scales. Section 7: Research and references: Summaries of normative, reliability, and validity studies and references to all sections. Oxford, England: Oxford Psychologists Press/San Antonio, TX: The Psychological Corporation.

12) Cummins C, Seale M, Macente A, Certini D, Mastropaolo E, et al. 2018. A separated vortex ring underlies the flight of the dandelion. Nature, 562(7727):414–418. doi: 10.1038/s41586-018-0604-2.

13) DeVries GA, Brunnbauer HY, Jackson AM, Long B, Neltner BT, et al. 2007. Divalent metal nanoparticles. Science, 315 (5810): 358–361.

14) Eisenberg M, Guy R. 1979. A Proof of the Hairy Ball Theorem. The American Mathematical Monthly, 86 (7): 571–574. doi: 10.2307/2320587.

15) Fingelkurts AA, Fingelkurts, A.A. 2004. Making complexity simpler: Multivariability and metastability in thebrain. Int. J.Neurosci. 2004, 114, 843–862; doi: 10.1080/00207450490450046.

16) Fingelkurts AA. Fingelkurts AA. 2015. Operational Architectonics methodology for EEG analysis: Theory and results. Neuromethods, 91: 1–59, doi: 10.1007/7657_2013_60.

17) Fingelkurts AA, Fingelkurts AA. 2017. Information flow in the brain: Ordered sequences of metastable states. Information, 8, 22; doi: 10.3390/info8010022.

18) Gersner R, Oberman L, Sanchez MJ, Chiriboga N, Kaye HL, et al. 2016. H-coil repetitive transcranial magnetic stimulation for treatment of temporal lobe epilepsy: A case report. Epilepsy Behav Case Rep. 2016; 5: 52–56. doi: 10.1016/j.ebcr.2016.03.001.

19) Gies H, Lippoldt St. 2015. Global surpluses of spin-base invariant fermions, Physics Letters B, 743: 415–419.

20) Jan MM, Alsallum MS, Sadler MR. 2017. Transcranial Magnetic Stimulation and Epilepsy. International Journal of Medical Science and Clinical Inventions 4(10): 3256–3260. DOI: 10.18535/ijmsci/v4i10.07 ICV 2015: 52.82.

21) Jaušovec N, Jaušovec K. 2010. Emotional Intelligence and Gender: A Neurophysiological Perspective. In A. Gruszka, G. Matthews, & B. Szymura (Eds.), Handbook of Individual Differences in Cognition (pp. 109–126). New York, NY: Springer New York. Retrieved from http://www.springerlink.com/index/10.1007/978-1-4419-1210-7_7.

22) Kohli S, Casson A. 2019. Machine learning validation of EEG+tACS artefact removal. Journal of Neural Engineering. https://doi.org/10.1088/1741-2552/ab58a3.

23) Lang PJ, Bradley MM, Cuthbert BN. 2005. International affective picture system (IAPS): Affective ratings of pictures and instructional manual. Technical Report A-6. Bainesville, FL. University of Florida.

24) Layer M, Forgan EM. 2010. Magnetic flux lines in type-II superconductors and the “hairy ball theorem”. Nature Comm. 1, 45.

25) Li J Ha NS, Liu TL, van Dam RM, Kim C-J. 2019. Ionic-surfactant-mediated electro-dewetting for digital microfluidics. Nature 572, pages507–510.

26) Liang L, Hum SV. 2013. A Low-Profile Antenna With Quasi-Isotropic Pattern for UHF RFID Applications, IEEE Antennas and Wireless Propagation Letters, 12, 210 – 213.

27) Milnor J. 1978. Analytic proofs of the “hairy ball”theorem, and the Brouwerfixed point theorem, American Mathematical Monthly, 1978;85 (7):521–524.

28) Miri M-A, Alù A. 2019. Exceptional points in optics and photonics. Science, 363 (6422), eaar7709. DOI: 10.1126/science.aar7709.

29) Modes CD, Warner M. 2012. Responsive nematic solid shells: Topology, compatibility and shape”, EPL, 2012; 97: 36007.

30) Sharmaa A, Reiterb G. 1996. Instability of Thin Polymer Films on Coated Substrates: Rupture, Dewetting, and Drop Formation. JColloid Interface Sci 178: 383–399.

31) Tao T; Tian, G. 2004. A singularity removal theorem for Yang-Mills fields in higher dimensions. J. Amer. Math. Soc. 17 (3): 557–593.

32) Tozzi A, Peters JF, Fingelkurts AA, Fingelkurts AA, Marijuán PC. 2017. Topodynamics of metastable brains. Physics of Life Reviews, 21, 1–20. http://dx.doi.org/10.1016/j.plrev.2017.03.001.

33) Tozzi A, Papo D. 2019. Projective mechanisms subtending real world phenomena wipe away cause effect relationships. Progress in Biophysics and Molecular Biology. DOI: 10.1016/j.pbiomolbio.2019.12.002.

34) Uhlenbeck K. 1982. Connections with L^p^ bounds on curvature. Comm. Math. Phys. 83 (1): 31–42.

35) Yao Z. 2019. Command of Collective Dynamics by Topological Defects in Spherical Crystals, Phys. Rev. Lett. 122, 228002.

